# JNJ-55511118 Stabilizes a Desensitized-like Conformation of γ8-Containing AMPA Receptors through Long-Range Allosteric Modulation

**DOI:** 10.64898/2025.12.10.693459

**Authors:** Paola Carrillo Flores, Vladimir Berka, Vasanthi Jayaraman

## Abstract

AMPA receptors (AMPARs), glutamate-gated ion channels, are dynamically regulated by auxiliary proteins such as TARP γ8. JNJ-55511118 (JNJ-118) is a selective inhibitor of γ8-containing AMPARs. The resting-state structure shows JNJ-118 binding near the transmembrane entrance at the AMPAR–γ8 interface, producing only localized structural effects. However, the mechanism by which JNJ-118 inhibits agonist-bound receptors remains unclear, as no activated-state structures are available. Using single-molecule fluorescence resonance energy transfer, fluorescence lifetime imaging, and electrophysiology, we examined how JNJ-118 reshapes AMPAR conformations in the presence and absence of glutamate. JNJ-118 shifts the glutamate-bound γ8-containing receptor toward a desensitized-like conformation by destabilizing the ligand-binding domain dimer interface, and this effect is supported by functional evidence showing allosteric competition between cyclothiazide and JNJ-118. Although the dimer interface adopts a configuration similar to that of AMPARs lacking TARPs, fluorescence lifetime measurements confirm that γ8 remains associated with the receptor. These findings demonstrate that under agonist-bound conditions, JNJ-118 acts as a long-range allosteric modulator that stabilizes a desensitized-like inhibited state without disrupting AMPAR–γ8 complex integrity.

## Introduction

α-Amino-3-hydroxy-5-methyl-4-isoxazolepropionic acid receptors (AMPARs), a subtype of glutamate-gated ion channels, mediate fast excitatory neurotransmission in the central nervous system and are essential for synaptic plasticity, learning, and memory (Hansen et al., 2021). Dysregulation of AMPAR activity has been implicated in neurological and psychiatric conditions, including epilepsy, chronic pain, depression, and neurodegenerative disorders (Hansen et al., 2021). AMPARs are dynamically modulated by auxiliary proteins, including transmembrane AMPAR regulatory proteins (TARPs) (Fukata et al., 2005; Milstein and Nicoll, 2008; Hansen et al., 2021). Among these, TARP γ8 plays an important role in the forebrain and hippocampus, regions associated with learning and memory (Rouach et al., 2005; Fukaya et al., 2006). γ8 alters AMPAR gating properties and pharmacology, with AMPARs showing resensitization under certain conditions, a phenomenon in which receptors return to an open state during sustained glutamate exposure (Kato et al., 2010a; Kato et al., 2010b; Riva et al., 2017; Carrillo et al., 2020).

Recently, highly selective allosteric modulators such as JNJ-55511118 (JNJ-118) have been developed, showing >1000-fold preference for γ8-containing AMPARs and no effect on isolated AMPARs or those associated with other auxiliary proteins (Gardinier et al., 2016; Kato et al., 2016; Maher et al., 2016; Hansen et al., 2021). Given this selectivity, these modulators have the potential to reduce pathological γ8-containing AMPAR activity with minimal off-target effects (Gardinier et al., 2016; Hoffman et al., 2021). The functional effects of JNJ-118 are well characterized (Gardinier et al., 2016; Dohrke et al., 2020; Coombs et al., 2022). It inhibits both peak and steady-state glutamate-evoked currents, decreases channel conductance, and accelerates desensitization (Coombs et al., 2022). In comparison, these electrophysiological studies provide detailed insights into functional modulation, the structural and dynamic basis for its mechanism remains poorly understood.

Recent cryo-electron microscopy (cryo-EM), mutagenesis, and molecular dynamics studies have provided structural insights into γ8-containing AMPARs bound to negative allosteric modulators, including JNJ-118, in the resting state (Dohrke et al., 2020; Zhang et al., 2023). These structures show that JNJ-118 binds at the AMPAR–γ8 interface within a lipid-accessible pocket near the extracellular end of the transmembrane segments, reorganizing the transmembrane helices and channel gate to stabilize the closed channel. However, these structures represent the resting state in the absence of agonists, and structural data for glutamate-bound AMPARs in complex with JNJ-118 are lacking. Thus, the mechanism of inhibition in the active state remains unclear.

Here, we used single-molecule fluorescence resonance energy transfer (smFRET) to investigate the conformational dynamics of the GluA2 subunit of AMPARs in complex with TARP γ8, in the presence and absence of JNJ-118, under both resting and active conditions. Our studies focused on the dimer interface within the ligand-binding domain (LBD), a region previously shown to be critical for allosteric inhibition and desensitization (Sun et al., 2002; Gonzalez et al., 2010; Dürr et al., 2014; Yelshanskaya et al., 2014; Hale et al., 2024).

GYKI, an allosteric inhibitor of AMPARs that binds at the entrance to the transmembrane segments, induces localized conformational changes near its binding site in the resting state (Yelshanskaya et al., 2016), but shows long-range conformational changes at the LBD dimer interface when studied in the presence of the agonist glutamate (Hale et al., 2024). This indicates that GYKI inhibition of the active state of the receptor occurs through stabilization of a decoupled configuration characteristic of the desensitized state (Hale et al., 2024). We aimed to determine whether such a mechanism is universal and also applies to modulators such as JNJ-118. Thus, we focused our smFRET analysis targeting the dimer interface of the LBD.

To complement smFRET, we performed electrophysiology using cyclothiazide, which binds to the dimer interface and stabilizes a coupled dimer, promoting the open channel state. This provides an indirect method to test allosteric communication between JNJ-118 binding near the transmembrane and cyclothiazide binding at the dimer interface in cells. We also used fluorescence lifetime imaging (FLIM) of proteins expressed in HEK-293 cells, by labeling AMPA receptor with acceptor and TARP γ8 with donor fluorophores, to determine if there were any large-scale distance changes between the extracellular domains of TARP γ8 and AMPA receptors upon binding JNJ-118.

Our combined fluorescence and electrophysiological studies show that JNJ-118 inhibits AMPARs in the presence of glutamate by stabilizing a desensitized-like conformation, with the dimer interface at the LBD being decoupled. The changes induced by JNJ-118 are, however localized, and there are no large-scale distance changes observed between the extracellular domains of TARP γ8 and AMPA receptors.

## Materials and methods

### Cloning and mutagenesis

Wild-type Rattus norvegicus GluA2-flip (unedited Q isoform) was mutated such that all extracellular nondisulfide-bonded cysteines were changed to serines (89, 196, and 436). For the smFRET measurement of distances within the dimer, a cysteine was introduced at position 467 to measure the distance between the proximal A and D subunits at the LBD. For measurement of intersubunit distance changes in the ATD of the AMPA receptor, a cysteine at position 23 was introduced in the cysteine-light GluA2 construct. This construct allows measurement across the dimers between the proximal B and D subunits at the ATD. To investigate the effect of auxiliary protein γ8 on the AMPA receptor, we generated tandem constructs of the cysteine-light GluA2 mutants with γ8 in which the carboxy-terminus of GluA2 was appended to the amino-terminus of γ8 by a Gly–Ser linker using Gibson assembly. For measuring AMPA-γ8 distances using FLIM, Amber TAG stop codons were introduced at position 73 of γ8, and the original stop codon was mutated to TAA. To label surface GluA2, a CLIP-tag was inserted within the ATD-LBD linker of GluA2 at position 371 aa. All mutagenesis was performed using standard site-directed mutagenesis procedures, and the resulting constructs were analyzed by sequencing. The functionality of the FRET constructs has been previously established (Carrillo et al., 2020).

### smFRET measurements

HEK293T cells were transfected using the manufacturer’s protocol for jetPRIME (Polyplus-transfection) with 10 µg of DNA and grown in media containing 30 µM NBQX. The cells were harvested after 24 h and washed with extracellular buffer (160 mM NaCl, 1.8 mM MgCl2, 1 mM CaCl2, 3 mM KCl, 10 mM glucose, and 10 mM HEPES, pH 7.4). The cells were then labeled with 300 nM each of maleimide derivatives of Alexa Fluor 555 donor and Alexa Fluor 647 acceptor fluorophores (Invitrogen). After being washed to remove excess fluorophores, the cells were solubilized by nutating in the dark for 1 h in chilled PBS containing 10 mM lauryl maltose neopentyl glycol and 0.25 mM cholesteryl hemisuccinate with protease inhibitor (Thermo Fisher Scientific). The lysed cells were centrifuged at 100,000×g at 4°C for 1 h, and the supernatant was used for smFRET sample preparation. Slides for smFRET studies were prepared and measurements taken as described previously (Litwin et al., 2019; MacLean et al., 2019; Bhatia et al., 2020; Carrillo et al., 2020; Durham et al., 2020a; Durham et al., 2020b; Litwin et al., 2020). A 1-ms resolution was used to acquire the photon counts produced per donor and acceptor excitation, then binned to 5 ms and denoised using wavelet decomposition (Taylor et al., 2010; Taylor and Landes, 2011). The background-corrected signal was used to calculate the FRET efficiency. Step transition and state identification analysis were used to determine the optimal number of states for the distribution of FRET efficiencies found in the obtained FRET data (Shuang et al., 2014). Observed efficiency histograms were fit to Gaussian distributions. Distance was determined from the FRET efficiencies using the Förster equation:

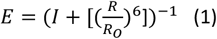

where R is the distance between the dyes, and R0 is the distance at half-maximal efficiency. The R_0_ is 51 Å for the Alexa Fluor 555–Alexa Fluor 647 fluorophore pair used for these experiments.

### Electrophysiology

We transfected HEK293T cells using iMFectin (GenDEPOT) following the manufacturer’s instructions with GluA2/γ8 tandem construct, and enhanced GFP DNA at a mass ratio of 1:0.2 µg per 2 ml of media. For whole-cell recordings, cells were replated after 4–6 h at a low density in fresh media containing 30 µM NBQX. Whole-cell patch-clamp recordings were performed 24 h after transfection using 3–5 MΩ resistance fire-polished borosilicate glass pipettes filled with the following internal solution: 135 mM CsF, 33 mM CsCl, 2 mM MgCl2, 1 mM CaCl2, 11 mM EGTA, and 10 mM HEPES, pH 7.4. The external solution consisted of 150 mM NaCl, 1 mM CaCl2, and 10 mM HEPES, pH 7.4.

Solutions with no added ligand, 10 mM glutamate, 10 mM glutamate with JNJ-118, 10 mM glutamate with 30 µM CTZ, and 10 mM glutamate with 30 µM CTZ in the presence of JNJ-118 were locally applied to lifted cells using a stepper motor system (SF-77B; Warner Instruments) with triple-barrel tubing.

Recordings were performed using an Axopatch 200B amplifier (Molecular Devices) at −60-mV hold potential, acquired at 10 kHz using pCLAMP10 software (Molecular Devices), and filtered online at 5 kHz.

### Fluorescence lifetime imaging

HEK-293T cells were plated on poly-d-lysine–coated glass coverslips and transfected 24 hours later with GluA2-CLIP and γ8-73. To incorporate the unnatural amino acid p-acetyl-l-phenylalanine into the γ8 protein during translation, cells were co-transfected with plasmids containing suppressor tRNA_CUA_ and the p-acetyl-l-phenylalanyl-tRNA synthetase, along with the GluA2 and γ8 plasmids. After transfection, the medium was supplemented with 500 μM amino acid p-acetyl-l-phenylalanine (AcF) (RSP Amino Acids). To measure FLIM, GluA2-CLIP was labeled with photostable fluorescent substrates CLIP-Surface 649 (New England Biolabs) and Alexa 555 hydrazide (Thermo Fisher Scientific). The experiments were performed after 24-48 hours of transfection. The cells were washed three times with an extracellular buffer solution (145 mM NaCl, 1 mM CaCl2, 3 mM KCl, 10 mM HEPES, and 10 mM glucose), incubated with 5 μM Clip Surface and 1.5 μM of Alexa 555 hydrazide, and then placed back in the incubator for a 2-hour period. After incubation, the cells were washed three times with extracellular buffer solution, and imaging experiments were performed.

For FLIM imaging, a MicroTime 200 confocal fluorescence microscope (PicoQuant, Berlin, Germany) was used, consisting of an inverted microscope (IX73, Olympus) equipped with an Olympus PlanApo 100×/numerical aperture 1.4 oil immersion objective. Fluorescence recordings were performed in the time-correlated single-photon counting mode using single-photon sensitive detectors (PicoQuant, Berlin, Germany). A 530-nm pulsed laser diode was used for excitation, band-pass filter of 582/64 nm was used to measure donor emission, and band-pass filter of 690/70 nm was used to measure acceptor emission. Data acquisition and analysis were performed by the SymPhoTime 64 software version 2.4 (PicoQuant, Berlin, Germany). The photons from the plasma membranes of the cells were selected to obtain the fluorescence intensity decay. The fluorescence intensity decays were analyzed using SigmaPlot. Data from 3 days were used to obtain mean and SE values for the lifetimes.

### Statistics

All electrophysiological and imaging data were statistically analyzed with the Student’s paired t test, with significance taken as P < 0.05. These tests were performed using GraphPad Prism software (GraphPad Software).

For smFRET, data were analyzed using Origin 9.0 (OriginLab Corp.), MATLAB (MathWorks), and Excel (Microsoft Corp.). At least three experiments were conducted for each condition. The numbers of molecules used for the histograms for L467C glutamate, L467C/γ8 apo, L467C/γ8 JNJ-118, L467C/γ8 glutamate, L467C/γ8 glutamate + JNJ-118, D23C/γ8 glutamate, D23C/γ8 glutamate + JNJ-118 were 90, 86, 91, 138, 97, 98 and 98, respectively.

## Results

### Effect of JNJ-118 on the conformational landscape of the LBD Dimer Interface of AMPA Receptors in Complex with TARP γ8

To investigate the conformational effects of JNJ-118 on γ8-containing AMPARs, we chose sites that are known to show large-scale conformational changes, the D1-D1 dimer interface at the LBD and the intersubunit interface at the ATD (Figure 1). For performing smFRET measurements at the LBD dimer interface, we used a cysteine-light GluA2 construct with an L467C mutation (Shaikh et al., 2016; Carrillo et al., 2020; Hale et al., 2024), which enables specific donor and acceptor fluorophore labeling at this site. This site reflects dimer decoupling during desensitization, with Cα distances of 31– 34 Å in the resting state and 40–42 Å in the desensitized state (Figure 1). These distances are well suited for Alexa555-Alexa647 donor-acceptor fluorophores (R_0_ = 51 Å), with the expected FRET efficiencies for these distances of 0.95–0.92 and 0.81–0.76 within the dimer in the resting and desensitized states. The FRET efficiencies expected for the distance across the dimer (Cα distances of 60–66 Å), are in the 0.27– 0.18 range, thus ensuring that the measurements would reflect the changes within the dimer without interference from across the dimer. To study distances across the amino-terminal domain (ATD), we chose site 23, which shows a distance of 50 Å in the intersubunit distance and a distance of 35 Å within the dimer. These distances translate to FRET efficiencies of 0.95-0.93 and 0.79-0.73, therefore allowing for selective investigation of the distance across the dimers.

**Figure 1.**
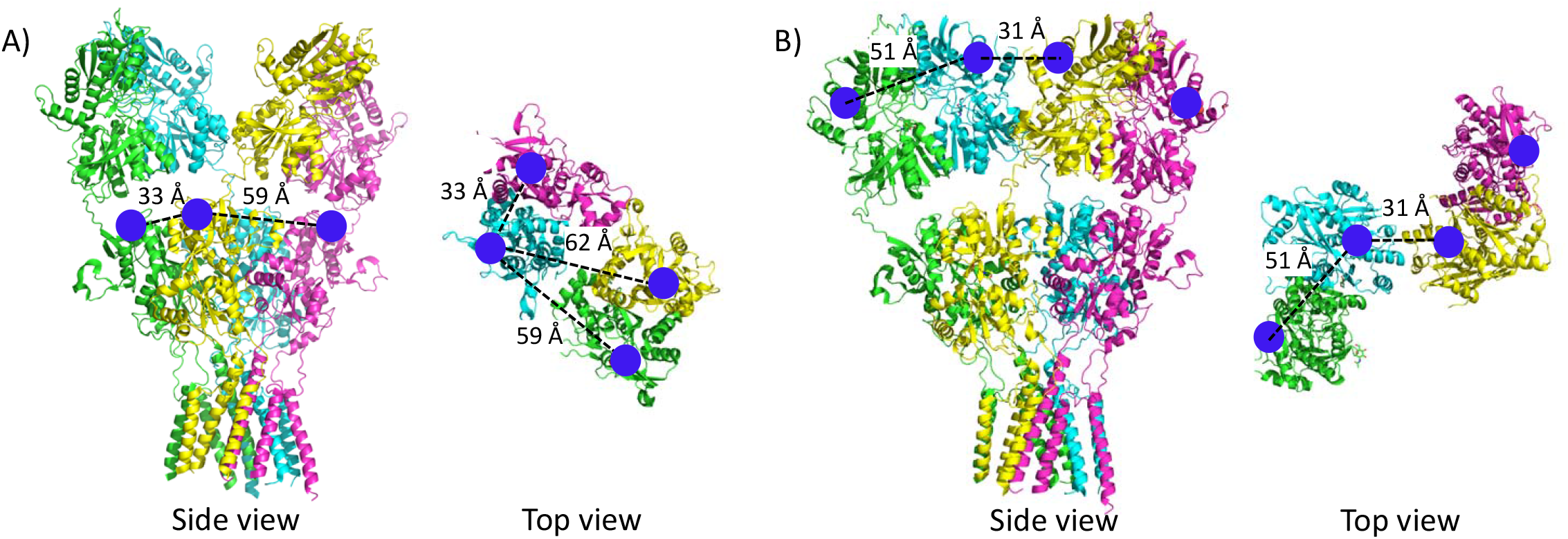
Labeling sites of smFRET constructs. A) Structure of the GluA2 receptor (PDB ID: 3KG2) with spheres indicating the fluorophore attachment site at the ligand-binding domain (L467). Both side and top views of the full-length apo GluA2 receptor are shown to illustrate the position of the labeling site. B) Structure of the GluA2 receptor (PDB ID: 3KG2) with spheres marking the fluorophore attachment site at the amino-terminal domain (D23). Side and top views of the full-length receptor highlight the relative orientation of the labeling site within the tetrameric assembly.

Using the LBD construct, smFRET measurements were performed under apo conditions for γ8-containing AMPARs, with 100 μM JNJ-118. The data showed two states at 0.92 (34 Å) and 0.72 (44 Å) (Figure 2a and Figure 2b), with interconversion within single traces, indicating conformational fluctuations within the same molecule. These distances are in the range expected for interdimer distance based on prior X-ray and cryo-EM data for AMPARs. The absence of a low-FRET state corresponding to the distance across the dimers suggests that this distance is longer and beyond the range probed by the Alexa555-Alexa647 donor–acceptor pair. The smFRET landscape for the apo state of γ8-containing AMPARs with JNJ-118 is similar to the smFRET for the apo state of γ8-containing AMPARs (without JNJ-118) (Figure 2c). Thus, JNJ-118 binding does not affect the LBD dimer interface in the apo state. This observation is consistent with cryo-EM data showing localized changes near the binding site and transmembrane segments, but no changes at the LBD dimer interface (Zhang et al., 2023).

**Figure 2.**
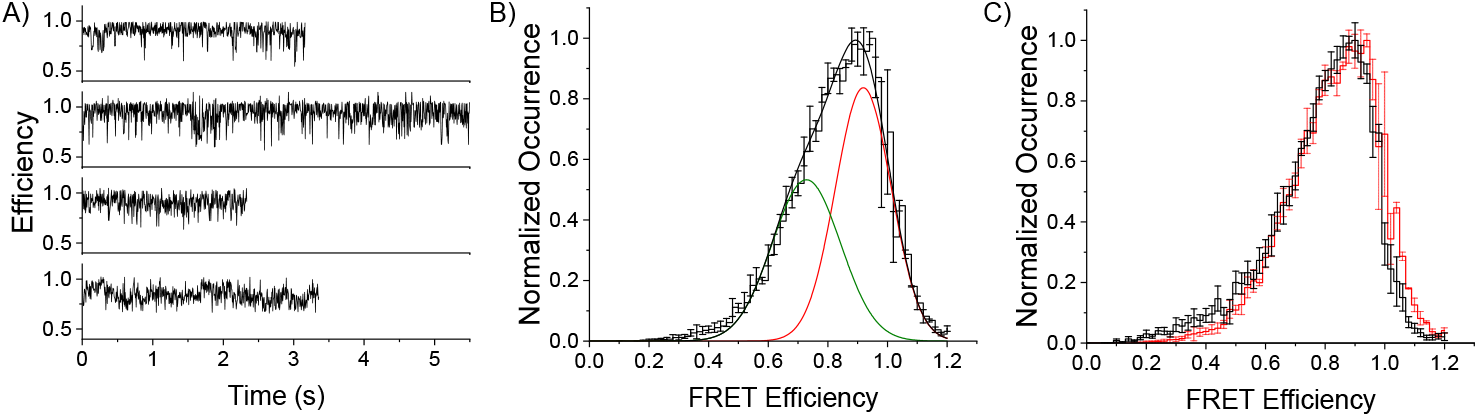
Conformational landscape at the LBD of the GluA2/γ8 receptors. A) Representative FRET-efficiency traces for GluA2/γ8 receptors in the apo condition in the presence of 100 μM JNJ-118. B) smFRET histograms of GluA2/γ8 receptors under apo conditions in the presence of 100 μM JNJ-118, fitted with two Gaussian components. C) Comparison between cumulative smFRET efficiency histograms for the apo condition of GluA2/γ8 receptors in the absence (red) and in the presence (black) of JNJ-118.

In the presence of agonist (1 mM glutamate), however, the addition of 100 μM JNJ-118 significantly shifted the smFRET landscape for the LBD dimer interface of γ8-containing AMPARs (Figure 3a) to lower FRET efficiencies. Based on the individual smFRET trajectories (examples provided in Figure 3b) the smFRET histogram for the γ8-containing AMPARs in the presence of JNJ-118 could be fit to three FRET levels (0.88, 0.75, and 0.6) (Figure 3c). We have previously published data for the glutamate-bound form of γ8-containing AMPARs based on 51 molecules (Carrillo et al., 2020). In the present study, we expanded this dataset to include an additional 87 molecules, and the results remain similar to our earlier findings. This shift in population toward lower FRET efficiencies indicates increased separation at the D1–D1 interface upon JNJ-118 binding in the activated receptor complex.

**Figure 3.**
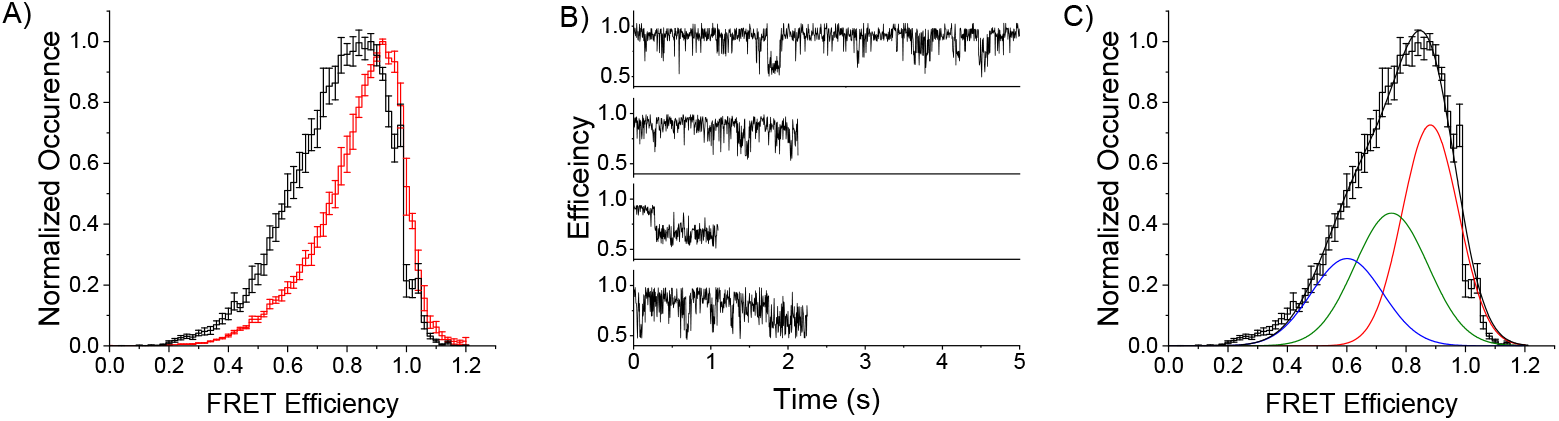
Conformational landscape at the LBD of GluA2/γ8 receptors in the active state. A) Comparison between cumulative smFRET efficiency histograms of GluA2/γ8 receptors in the presence of 1 mM glutamate, in the presence (black) and absence of 100 μM JNJ-118 (red). B) Representative FRET-efficiency traces for GluA2/γ8 receptors in the presence of 1 mM glutamate and 100 μM JNJ-118. C) smFRET histograms display the population distribution of GluA2/γ8 receptors under glutamate conditions, fitted with three Gaussian components.

Given the large changes observed at the D1-D1 interface upon the addition of 100 μM JNJ-118 in the glutamate bound form of γ8-containing AMPARs we used smFRET to determine if this was also observed at the ATD inter-dimer interface. The smFRET data show a shift in the FRET landscape upon addition of JNJ-118 to lower FRET states (Figure 4a). Based on the smFRET trajectories obtained from γ8-containing AMPARs in the presence of JNJ-118 (representative traces shown in Figure 4b) the histogram was fit to three distinct FRET levels (0.92, 0.72, and 0.48) (Figure 4c). These data suggest that there is decoupling observed in the ATD upon addition of JNJ-118, indicating that JNJ-118 has long-range effects on the AMPARs. However, it should be noted that the decoupling is significantly smaller than what we had previously seen for AMPARs alone in the absence of TARPs (Carrillo et al., 2020).

**Figure 4.**
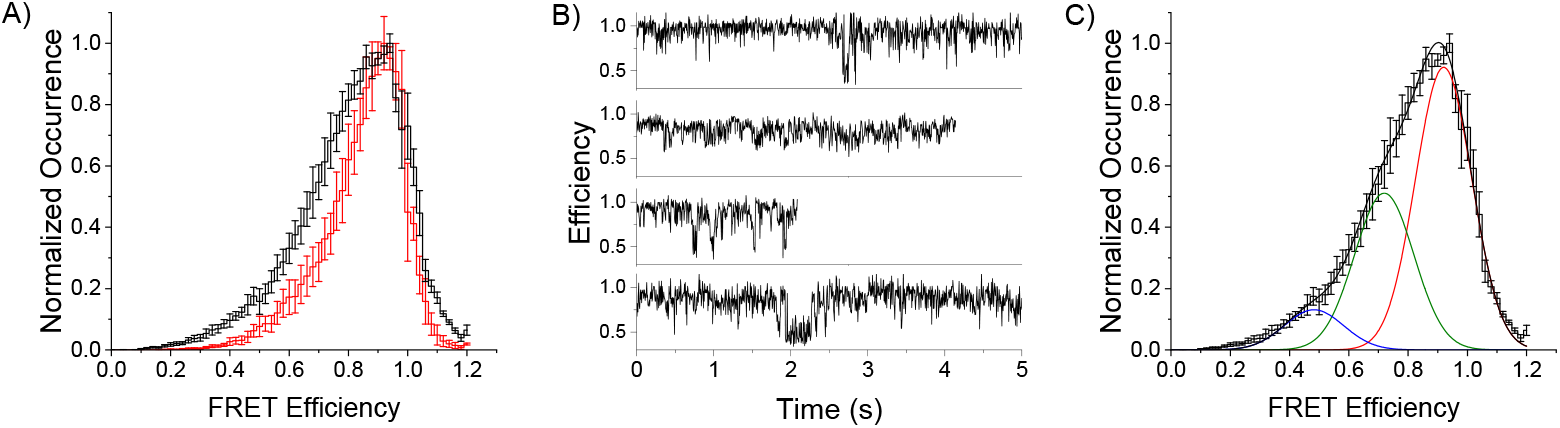
Conformational landscape at the ATD of GluA2/γ8 receptors in the active state. A) Comparison between cumulative smFRET efficiency histograms of GluA2/γ8 receptors in the presence of 1 mM glutamate, in the presence (black) and absence of 100 μM JNJ-118 (red). B) Representative FRET-efficiency traces for GluA2/γ8 receptors in the presence of 1 mM glutamate and 100 μM JNJ-118. C) smFRET histograms display the population distribution of GluA2/γ8 receptors under glutamate conditions, fitted with three Gaussian components.

### Dimer Decoupling at the LBD Probed with Allosteric Competitive Studies Using Cyclothiazide

To further verify the loss of the LBD coupled state of the dimer interface, we probed the changes at the dimer interface indirectly by investigating the effect of JNJ-118 on the binding of cyclothiazide, a drug that stabilizes the D1–D1 interface of the LBD. Functional recordings show that the dose response for inhibition by JNJ-118 is significantly shifted to higher concentrations in the presence of cyclothiazide EC_50_ of 3.5 ± 0.5 µM to 78 ± 8.5 µM (Figure 5). This shift indicates that cyclothiazide counteracts the inhibitory effect of JNJ-118 by reinforcing structural coupling at the D1–D1 interface and establishes allosteric communication between the JNJ-118 binding site and dimer interface of the AMPAR LBD.

**Figure 5.**
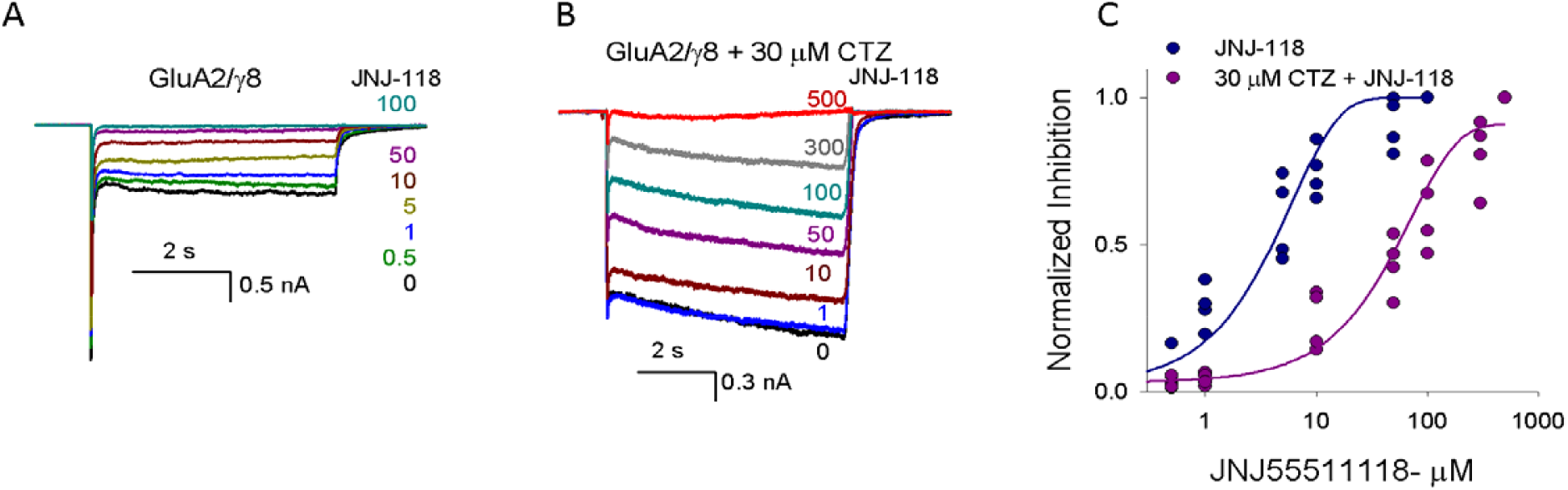
Reinforced D1–D1 coupling by cyclothiazide counteracts JNJ-118 inhibition. A) Representative whole-cell currents from GluA2/γ8 receptors activated by 10 mM glutamate in the presence of increasing concentrations of JNJ-118 (0, 0.5, 1, 5, 10, 50, 100 μM). B) Representative GluA2/γ8 currents evoked by 10 mM glutamate in the presence of 30 µM CTZ with varying concentrations of JNJ-118 (0, 1, 10, 50, 100, 300, 500 μM). C) The presence of 30 µM CTZ produces a shift in JNJ-118 potency, increasing the EC_50_ from 3.5 ± 0.5 µM to 78 ± 8.5 µM.

### Comparison of JNJ-118 bound form of γ8-containing AMPARs with desensitized AMPA receptors

Given that the dimer interface is decoupled for the glutamate-bound form of γ8-containing AMPARs in the presence of JNJ-118, indicating a desensitized-like state, we next investigated if this desensitized state is similar to the desensitized state observed in glutamate-bound AMPA receptors in the absence of TARPs. The smFRET histograms show that the conformational landscape of 100 μM JNJ-118-bound γ8-containing AMPARs in the presence of glutamate is similar to that of glutamate-bound AMPARs in the absence of TARP γ8 (Figure 6a). The similarity between these conditions suggests that binding of JNJ-118 releases the effect of TARP γ8 on the AMPA receptor LBD dimer interface. The data presented here for the glutamate-bound AMPARs in the absence of TARP γ8 is an expanded data set of previously published data (36 molecules) (Carrillo et al., 2020).

**Figure 6.**
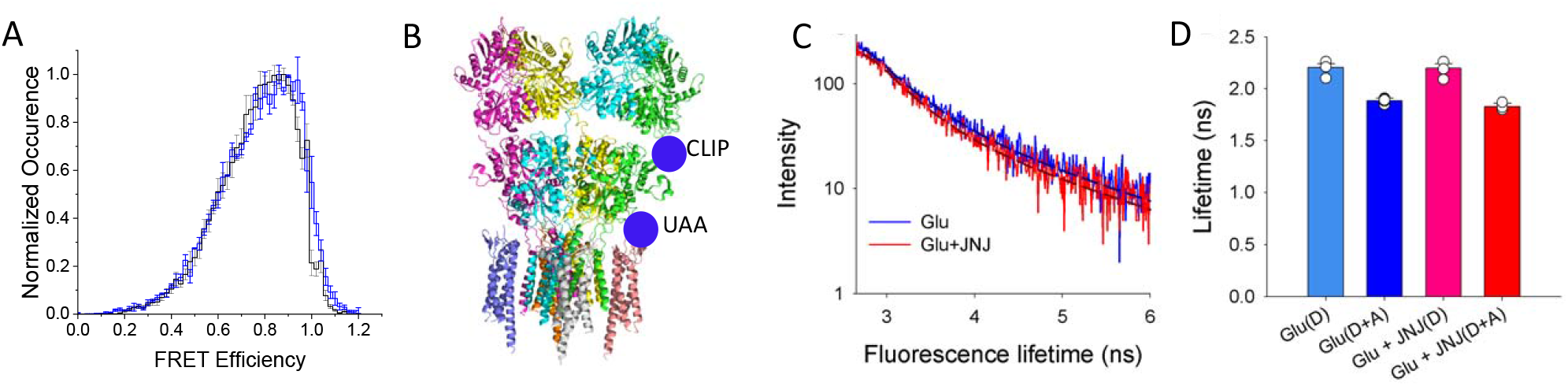
JNJ-118 promotes conformational states in γ8-AMPARs resembling desensitized AMPA receptors. A) In glutamate, JNJ-118–bound γ8-containing AMPARs display an smFRET landscape similar to desensitized AMPARs lacking TARP γ8 (blue). B) Sites for labeling for FLIM measurements using GluA2 and γ8. CLIP-tag was inserted within the ATD-LBD linker of GluA2 at position 371 and labeled with CLIP-Surface 649 (donor). p-acetyl-l-phenylalanine-tagged TARPs were labeled with Alexa 555 hydrazide (acceptor). C) Representative fluorescence lifetimes of donor- and acceptor-labeled cells expressing GluA2-CLIP and γ8 S73*. The lines show two exponential fits. D) Average Alexa 555 hydrazide lifetime in HEK293T cells co-expressing GluA2-CLIP and γ8 S73*, from at least three different days. Data represent mean lifetimes ± SEM, p = 0.075.

To determine if JNJ-118 binding dissociates TARP γ8 from AMPARs, we determined the distance between AMPARs and TARP γ8 using sites in the extracellular domain (Figure 6b). Fluorophores were tagged on γ8 by introducing the unnatural amino acid p-acetyl-l-phenylalanine at site 73 aa on the extracellular domain. The keto group of p-acetyl-l-phenylalanine could then be coupled to Alexa 555 hydrazide. To label surface GluA2, a CLIP-tag was inserted within the ATD-LBD linker of GluA2 at position 371 aa, coupled to Alexa Clip Surface 647 (Figure 6b). Fluorescence lifetime imaging microscopy (FLIM) was used to estimate the degree of FRET-based changes of the donor Alexa 555 hydrazide fluorescence lifetime. The donor only lifetime (γ8 S73*; Alexa 555 hydrazide) was 2.25 ± 0.04 ns and 2.2 ± 0.07, in the presence of glutamate and glutamate with 100 μM JNJ-118, respectively (Figure 6c). We observed a decrease in the fluorescence lifetime in the presence of donor and acceptor (Figure 6d), showing that there was FRET between the two fluorophores. The fluorescence lifetime for the FRET between the two sites was not significantly different between glutamate and glutamate with JNJ-118, showing that there were no significant distance changes between the two sites upon addition of JNJ-118. Thus, indicating that JNJ-118 does not cause a dissociation of the GluA2-γ8 complex. This is also consistent with the smFRET data at the ATD, which showed changes due to JNJ-118 binding, but the conformational landscape is not to the same extent as seen for AMPARs in the absence of TARPs.

## Discussion

Our study provides mechanistic insight into how the TARP γ8-selective negative allosteric modulator JNJ-118 inhibits γ8-containing AMPAR function. Using smFRET and electrophysiological assays with cyclothiazide, we show that JNJ-118 exerts state-dependent effects on the conformation of γ8-containing AMPARs. Specifically, JNJ-118 promotes destabilization of the D1–D1 interface of the ligand-binding domain in the glutamate-bound state, consistent with a desensitized-like conformation. In contrast, the apo-state conformation remains largely unaffected by JNJ-118 binding, in agreement with previous cryo-EM studies that showed only localized changes in the transmembrane domain in the absence of agonist (Zhang et al., 2023).

Our findings also clarify the extent to which JNJ-118 mimics or reverses TARP γ8 modulation. While JNJ-118 partially reduces the functional enhancements conferred by γ8, including decreased steady-state current, reduced conductance, and faster desensitization, it does not fully revert AMPAR gating to a TARPless profile (Coombs et al., 2022). This aligns with our FLIM observations, which shows no large-scale distance changes between the extracellular domains of AMPARs and γ8 upon addition of JNJ-118. Therefore, JNJ-118 can be viewed not as a pure antagonist of γ8 function, but as a modulator that selectively dampens TARP-enhanced gating while preserving complex integrity.

Further evidence for long-range allosteric effects comes from our functional interaction studies with cyclothiazide. Cyclothiazide is known to stabilize the D1–D1 interface (Sun et al., 2002) and enhance AMPAR activity by preventing desensitization (Yamada and Tang, 1993; Partin et al., 1994). Here, we show that cyclothiazide reduces the inhibitory efficacy of JNJ-118, as reflected by a rightward shift in the JNJ-118 dose–response curve in the presence of cyclothiazide. This functional antagonism supports a model of allosteric competition, in which positive and negative allosteric modulators act through distinct sites but converge on the same conformational pathway—the D1–D1 interface. This behavior closely parallels the action of other non-competitive AMPAR antagonists such as GYKI-52466, which also stabilize a desensitized-like conformation in agonist-bound receptors (Hale et al., 2024). This suggests that the allosteric pathway between the transmembrane segments and the dimer interface is bidirectional—not only important in mediating agonist-induced changes from the extracellular domain to the transmembrane domain, but also in communicating modulator binding from the transmembrane segment to the extracellular domain. This mechanism represents a general paradigm for the action of peripheral, TARP-dependent modulators and distinguishes their nuanced modulatory effects from the binary gating control exerted by pore blockers or orthosteric antagonists.

In conclusion, our findings demonstrate that JNJ-118 inhibits AMPAR activation by promoting D1–D1 interface decoupling in a state-dependent manner while preserving TARP association. This partial disruption of gating offers a mechanism for fine-tuned inhibition of AMPARs in a TARP-selective context. Our study also highlights the D1–D1 interface as a key conformational nexus for allosteric control and a promising target for future drug development.

## Acknowledgments

This project was supported by NIH grant R35-GM122528 (VJ). The authors used ChatGPT (OpenAI, version 5.1) for proofreading of the manuscript. The authors take full responsibility for the content.

## Author contributions

V.B. and V.J. designed research; P.C.F., and V.B. performed smFRET experiments and electrophysiological recordings; P.C.F., V.B. and V.J. analyzed data; and P.C.F., and V.J. wrote the paper.

## Data availability

The datasets generated during and/or analyzed during the current study are available from the corresponding author upon reasonable request.

## References

Bhatia, N.K., E. Carrillo, R.J. Durham, V. Berka, and V. Jayaraman. 2020. Allosteric Changes in the NMDA Receptor Associated with Calcium-Dependent Inactivation. Biophys J. 119:2349–2359.

Carrillo, E., S.A. Shaikh, V. Berka, R.J. Durham, D.B. Litwin, G. Lee, D.M. MacLean, L.M. Nowak, and V. Jayaraman. 2020. Mechanism of modulation of AMPA receptors by TARP-γ8. J Gen Physiol. 152.

Coombs, I.D., C.A. Sexton, S.G. Cull-Candy, and M. Farrant. 2022. Influence of the TARP gamma 8-Selective Negative Allosteric Modulator JNJ-55511118 on AMPA Receptor Gating and Channel Conductance. Molecular Pharmacology. 101:343–356.

Dohrke, J.-N., J.F. Watson, K. Birchall, and I.H. Greger. 2020. Characterizing the binding and function of TARP γ8-selective AMPA receptor modulators. Journal of Biological Chemistry. 295:14565–14577.

Durham, R.J., D.R. Latham, H. Sanabria, and V. Jayaraman. 2020a. Structural Dynamics of Glutamate Signaling Systems by smFRET. Biophys J. 119:1929–1936.

Durham, R.J., N. Paudyal, E. Carrillo, N.K. Bhatia, D.M. Maclean, V. Berka, D.M. Dolino, A.A. Gorfe, and V. Jayaraman. 2020b. Conformational spread and dynamics in allostery of NMDA receptors. Proc Natl Acad Sci U S A. 117:3839–3847.

Dürr, K.L., L. Chen, R.A. Stein, R. De Zorzi, I.M. Folea, T. Walz, H.S. McHaourab, and E. Gouaux. 2014. Structure and dynamics of AMPA receptor GluA2 in resting, pre-open, and desensitized states. Cell. 158:778–792.

Fukata, Y., A.V. Tzingounis, J.C. Trinidad, M. Fukata, A.L. Burlingame, R.A. Nicoll, and D.S. Bredt. 2005. Molecular constituents of neuronal AMPA receptors. J Cell Biol. 169:399–404.

Fukaya, M., M. Tsujita, M. Yamazaki, E. Kushiya, M. Abe, K. Akashi, R. Natsume, M. Kano, H. Kamiya, M. Watanabe, and K. Sakimura. 2006. Abundant distribution of TARP gamma-8 in synaptic and extrasynaptic surface of hippocampal neurons and its major role in AMPA receptor expression on spines and dendrites. Eur J Neurosci. 24:2177–2190.

Gardinier, K.M., D.L. Gernert, W.J. Porter, J.K. Reel, P.L. Ornstein, P. Spinazze, F.C. Stevens, P. Hahn, S.P. Hollinshead, D. Mayhugh, J. Schkeryantz, A. Khilevich, O. De Frutos, S.D. Gleason, A.S. Kato, D. Luffer-Atlas, P.V. Desai, S. Swanson, K.D. Burris, C. Ding, B.A. Heinz, A.B. Need, V.N. Barth, G.A. Stephenson, B.A. Diseroad, T.A. Woods, H. Yu, D. Bredt, and J.M. Witkin. 2016. Discovery of the First α-Amino-3-hydroxy-5-methyl-4-isoxazolepropionic Acid (AMPA) Receptor Antagonist Dependent upon Transmembrane AMPA Receptor Regulatory Protein (TARP) γ-8. J Med Chem. 59:4753–4768.

Gonzalez, J., M. Du, K. Parameshwaran, V. Suppiramaniam, and V. Jayaraman. 2010. Role of dimer interface in activation and desensitization in AMPA receptors. Proc Natl Acad Sci U S A. 107:9891–9896.

Hale, W.D., A. Montano Romero, C.U. Gonzalez, V. Jayaraman, A.Y. Lau, R.L. Huganir, and E.C. Twomey. 2024. Allosteric competition and inhibition in AMPA receptors. Nat Struct Mol Biol. 31:1669–1679.

Hansen, K.B., L.P. Wollmuth, D. Bowie, H. Furukawa, F.S. Menniti, A.I. Sobolevsky, G.T. Swanson, S.A. Swanger, I.H. Greger, T. Nakagawa, C.J. McBain, V. Jayaraman, C.M. Low, M.L. Dell’Acqua, J.S. Diamond, C.R. Camp, R.E. Perszyk, H. Yuan, and S.F. Traynelis. 2021. Structure, Function, and Pharmacology of Glutamate Receptor Ion Channels. Pharmacol Rev. 73:298–487.

Hoffman, J.L., S. Faccidomo, B.L. Saunders, S.M. Taylor, M. Kim, and C.W. Hodge. 2021. Inhibition of AMPA receptors (AMPARs) containing transmembrane AMPAR regulatory protein γ-8 with JNJ-55511118 shows preclinical efficacy in reducing chronic repetitive alcohol self-administration. Alcohol Clin Exp Res. 45:1424–1435.

Kato, A.S., K.D. Burris, K.M. Gardinier, D.L. Gernert, W.J. Porter, J. Reel, C. Ding, Y. Tu, D.A. Schober, M.R. Lee, B.A. Heinz, T.E. Fitch, S.D. Gleason, J.T. Catlow, H. Yu, S.M. Fitzjohn, F. Pasqui, H. Wang, Y. Qian, E. Sher, R. Zwart, K.A. Wafford, K. Rasmussen, P.L. Ornstein, J.T. Isaac, E.S. Nisenbaum, D.S. Bredt, and J.M. Witkin. 2016. Forebrain-selective AMPA-receptor antagonism guided by TARP γ-8 as an antiepileptic mechanism. Nat Med. 22:1496–1501.

Kato, A.S., M.B. Gill, M.T. Ho, H. Yu, Y. Tu, E.R. Siuda, H. Wang, Y.W. Qian, E.S. Nisenbaum, S. Tomita, and D.S. Bredt. 2010a. Hippocampal AMPA receptor gating controlled by both TARP and cornichon proteins. Neuron. 68:1082–1096.

Kato, A.S., M.B. Gill, H. Yu, E.S. Nisenbaum, and D.S. Bredt. 2010b. TARPs differentially decorate AMPA receptors to specify neuropharmacology. Trends Neurosci. 33:241–248.

Litwin, D.B., R.J. Durham, and V. Jayaraman. 2019. Single-Molecule FRET Methods to Study Glutamate Receptors. Methods Mol Biol. 1941:3–16.

Litwin, D.B., N. Paudyal, E. Carrillo, V. Berka, and V. Jayaraman. 2020. The structural arrangement and dynamics of the heteromeric GluK2/GluK5 kainate receptor as determined by smFRET. Biochim Biophys Acta Biomembr. 1862:183001.

MacLean, D.M., R.J. Durham, and V. Jayaraman. 2019. Mapping the Conformational Landscape of Glutamate Receptors Using Single Molecule FRET. Trends Neurosci. 42:128–139.

Maher, M.P., N. Wu, S. Ravula, M.K. Ameriks, B.M. Savall, C. Liu, B. Lord, R.M. Wyatt, J.A. Matta, C. Dugovic, S. Yun, L. Ver Donck, T. Steckler, A.D. Wickenden, N.I. Carruthers, and T.W. Lovenberg. 2016. Discovery and Characterization of AMPA Receptor Modulators Selective for TARP-γ8. J Pharmacol Exp Ther. 357:394–414.

Milstein, A.D., and R.A. Nicoll. 2008. Regulation of AMPA receptor gating and pharmacology by TARP auxiliary subunits. Trends Pharmacol Sci. 29:333–339.

Partin, K.M., D.K. Patneau, and M.L. Mayer. 1994. Cyclothiazide differentially modulates desensitization of alpha-amino-3-hydroxy-5-methyl-4-isoxazolepropionic acid receptor splice variants. Mol Pharmacol. 46:129–138.

Riva, I., C. Eibl, R. Volkmer, A.L. Carbone, and A.J. Plested. 2017. Control of AMPA receptor activity by the extracellular loops of auxiliary proteins. Elife. 6.

Rouach, N., K. Byrd, R.S. Petralia, G.M. Elias, H. Adesnik, S. Tomita, S. Karimzadegan, C. Kealey, D.S. Bredt, and R.A. Nicoll. 2005. TARP gamma-8 controls hippocampal AMPA receptor number, distribution and synaptic plasticity. Nat Neurosci. 8:1525–1533.

Shaikh, S.A., D.M. Dolino, G. Lee, S. Chatterjee, D.M. MacLean, C. Flatebo, C.F. Landes, and V. Jayaraman. 2016. Stargazin Modulation of AMPA Receptors. Cell Rep. 17:328–335.

Shuang, B., D. Cooper, J.N. Taylor, L. Kisley, J. Chen, W. Wang, C.B. Li, T. Komatsuzaki, and C.F. Landes. 2014. Fast Step Transition and State Identification (STaSI) for Discrete Single-Molecule Data Analysis. J Phys Chem Lett. 5:3157–3161.

Sun, Y., R. Olson, M. Horning, N. Armstrong, M. Mayer, and E. Gouaux. 2002. Mechanism of glutamate receptor desensitization. Nature. 417:245–253.

Taylor, J.N., and C.F. Landes. 2011. Improved resolution of complex single-molecule FRET systems via wavelet shrinkage. J Phys Chem B. 115:1105–1114.

Taylor, J.N., D.E. Makarov, and C.F. Landes. 2010. Denoising single-molecule FRET trajectories with wavelets and Bayesian inference. Biophys J. 98:164–173.

Yamada, K.A., and C.M. Tang. 1993. Benzothiadiazides inhibit rapid glutamate receptor desensitization and enhance glutamatergic synaptic currents. J Neurosci. 13:3904–3915.

Yelshanskaya, M.V., M. Li, and A.I. Sobolevsky. 2014. Structure of an agonist-bound ionotropic glutamate receptor. Science. 345:1070–1074.

Yelshanskaya, M.V., A.K. Singh, J.M. Sampson, C. Narangoda, M. Kurnikova, and A.I. Sobolevsky. 2016. Structural Bases of Noncompetitive Inhibition of AMPA-Subtype Ionotropic Glutamate Receptors by Antiepileptic Drugs. Neuron. 91:1305–1315.

Zhang, D., R. Lape, S.A. Shaikh, B.K. Kohegyi, J.F. Watson, O. Cais, T. Nakagawa, and I.H. Greger. 2023. Modulatory mechanisms of TARP γ8-selective AMPA receptor therapeutics. Nature Communications. 14:1659.

